# Investigating the mechanisms of antibody binding to alpha-synuclein for the treatment of Parkinson’s Disease

**DOI:** 10.1101/2024.02.27.582350

**Authors:** Malcolm C. Harrison, Pin-Kuang Lai

## Abstract

Parkinson’s Disease (PD) is an idiopathic neurodegenerative disorder with the second-highest prevalence rate behind Alzheimer’s Disease. The pathophysiological hallmarks of PD are both degeneration of dopaminergic neurons in the substantia nigra pars compacta and the inclusion of misfolded alpha-synuclein (α-syn) aggregates known as Lewy bodies. Despite decades of research for potential PD treatments, none have been developed, and developing new therapeutic agents is a time-consuming and expensive process. Computational methods can be used to investigate the properties of drug candidates currently undergoing clinical trials to determine their theoretical efficiency at targeting α-syn. Monoclonal antibodies (mAbs) are biological drugs with high specificity, and Prasinezumab (PRX002) is a mAb currently in Phase II, which targets the C-terminus (AA 118-126) of α-syn. We utilized BioLuminate and PyMol for structure prediction and preparation of the fragment antigen-binding (Fab) region of PRX002 and 34 different conformations of α-syn. Protein-protein docking simulations were performed using PIPER, and 3 of the docking poses were selected based on the best fit. Molecular dynamics simulations were conducted on the docked protein structures for 1000 ns, and hydrogen bonds, electrostatic, and hydrophobic interactions were analyzed using MDAnalysis to determine which residues were interacting and how often. Hydrogen bonds were shown to form frequently between the HCDR2 region of PRX002 and α-syn. Free energy was calculated to determine binding affinity. The predicted binding affinity shows a strong antibody-antigen attraction between PRX002 and α-syn. RMSD was calculated to determine the conformational change of these regions throughout the simulation. The mAb’s developability was determined using computational screening methods. Our results demonstrate the efficiency and developability of this therapeutic agent.

## 1. Introduction

Parkinson’s Disease (PD) is an idiopathic neurodegenerative disease that affects 2-3% of the population over 65 [1]. The pathophysiological hallmarks of PD are the degeneration of dopaminergic neurons in the substantia nigra pars compacta (SNpc), as well as the inclusion of cytotoxic Lewy bodies [2]. The primary constituent of Lewy bodies is misfolded aggregates of alpha-synuclein (α-syn) [3], consisting of 140 amino acid residues encoded by the synuclein alpha (SNCA) gene, which upon mutation, causes familial PD [4]. The loss of dopaminergic neurons in the SNpc causes a decrease in dopamine (DA) levels in the corpus striatum, generating deregulation of the basal ganglia systems, leading to motor symptoms such as resting tremor, rigidity, bradykinesia, and loss of static stability [5]. Depletion of DA levels can reach 80% before any changes in tonic DA can be measured [6], showing the need to identify PD before the disease has caused irreparable damage. Siderowf et al. showed that α-syn seed amplification assays can be applied for the accurate identification of PD patients and those at high risk for PD [7].

The discovery, development, and manufacturing of new drug candidates is a long, laborious, and expensive process, with the average cost for research and development of a single drug candidate at $1.1 billion [8]. In the last 30 years, monoclonal antibodies (mAbs) have come into the biopharmaceutical landscape as therapeutic agents rather than tools for scientific analysis [9]. These medications are highly specific, as they are designed for a specific epitope and administered either intravenously or subcutaneously [10]. MAbs have been developed to treat oncological, cardiovascular, kidney, and neurological diseases [11]. The biggest challenge with developing mAbs for central nervous system disorders is the large size of biological drugs [12] attempting to pass through the blood-brain barrier (BBB) [13]. Recent successes in the development of mAbs for Alzheimer’s Disease (AD) show the promise of targeting and lowering the number of protein aggregates in the brain [14] [15].

Prasinezumab (PRX002) is a mAb developed by Roche and Prothena that targets the C-terminus (AA 118-126) of α-syn [16] [17]. PRX002 is currently in clinical trial stage 2b after a promising phase 1b trial, which showed its high affinity for peripheral α-syn as well as no fatalities, severe adverse effects, or anti-PRX002 antibodies formed. Compared to another anti-α-syn mAb, Cinpanemab, PRX002 recognizes and binds to the C-terminus of aggregated and monomeric α-syn better than the N-terminus binding counterpart, which Biogen is no longer developing [18].

In the past three decades, computer-aided drug discovery methods have been extremely influential in the biopharmaceutical industry [19]. The ability to speed up and reduce the price of developing therapeutics is an extremely attractive characteristic of utilizing computational tools in research and development settings. Of these technologies, molecular dynamics (MD) and machine learning (ML) have had the highest influence. MD can be used to simulate interactions between proteins and ligands [20], antibody-antigen binding [21], as well as discovering novel binding sites [22]. ML has great applications for utilizing the physicochemical properties of drug-like compounds to predict if a projected drug will be a successful candidate in both clinical trials and mass production [23] [24]. These tools have been extremely valuable in the search for a cure for challenging diseases, including neurological conditions such as PD [25, 26, 27] and AD [28].

The administration of mAbs presents a challenge for PD patients, as the cognitive and motor symptoms of the disease greatly affect driving ability and safety [29], leading to increased dependency on a caregiver for transportation. Subcutaneous (SC) administration of PRX002 would allow PD patients to safely administer this medication from their homes, potentially increasing safety and adherence. The challenge with SC administration is the large size of mAbs, which can lead to aggregation and elevated viscosity [30].

In this paper, we utilize computational methods to model the interactions between the fragment antigen binding (Fab) region of PRX002 with α-syn *in silico*, simulate binding between these two proteins, perform molecular dynamics (MD) simulations to analyze intermolecular forces taking place at the binding site, calculate the binding free energy of the antibody and antigen, determine the conformational changes of the complementarity determining regions (CDR) of the variable region (Fv) on both the heavy and light chain (V_H_ and V_L_ respectively), as well as test the potential for developability and similarity to clinical stage therapeutics. This mechanistic study will help evaluate the effectiveness of PRX002 and provide a workflow to optimize future drug discovery and development, reducing experimental costs and leading to more effective and tailored therapies.

## 2. Materials and Methods

### 2.1: Generation of Docked PRX002-α-syn Complex

The sequence data for PRX002 was obtained from the Kyoto Encyclopedia of Genes and Genomes (KEGG) [31], and Schrödinger software BioLuminate [32, 33, 34, 35, 36, 37] and PyMol [38] were used to predict the protein structure of the Fab region of the mAb, with Chothia numbering used to annotate the final models. Thirty-four different variations of α-syn were obtained from the Protein Data Bank (PDB ID 2KKW [34]), which all had different structures gathered by nuclear magnetic resonance imaging. Both proteins were prepared by removing water and adding missing side chain atoms using the Protein Preparation Wizard [39], and structure reliability reports were completed using BioLuminate. PIPER [40, 41] was used to perform protein-protein docking on PRX002 and all 34 variations of α-syn.

### 2.2: Simulation Preparation

Using Visual Molecular Dynamics (VMD, version 1.9.3) [42], the heavy chain and light chains of PRX002 and α-syn were separated, renumbered, and patches were used between cysteine residues on the Fab region to stabilize the disulfide bridges with CHARMM36m force field[]. The VMD plugin psfgen was then used to create a simulation-ready protein atom coordinate file (pdb) and protein structure file (psf). This structure then underwent minimization for 5000 steps using Nanoscale Molecular Dynamics (NAMD, version 2.14) [43]. Solvation and ionization were performed utilizing the solvate and autoionize VMD plugins using a PDB file generated by the minimization step, embedding the structure in a solvate box extending 10Å from the protein boundary (Figure 1). The CHARMM36m force field was utilized in the simulation along the three-point water model (TIP3P) [44, 45].

**Figure 1:**
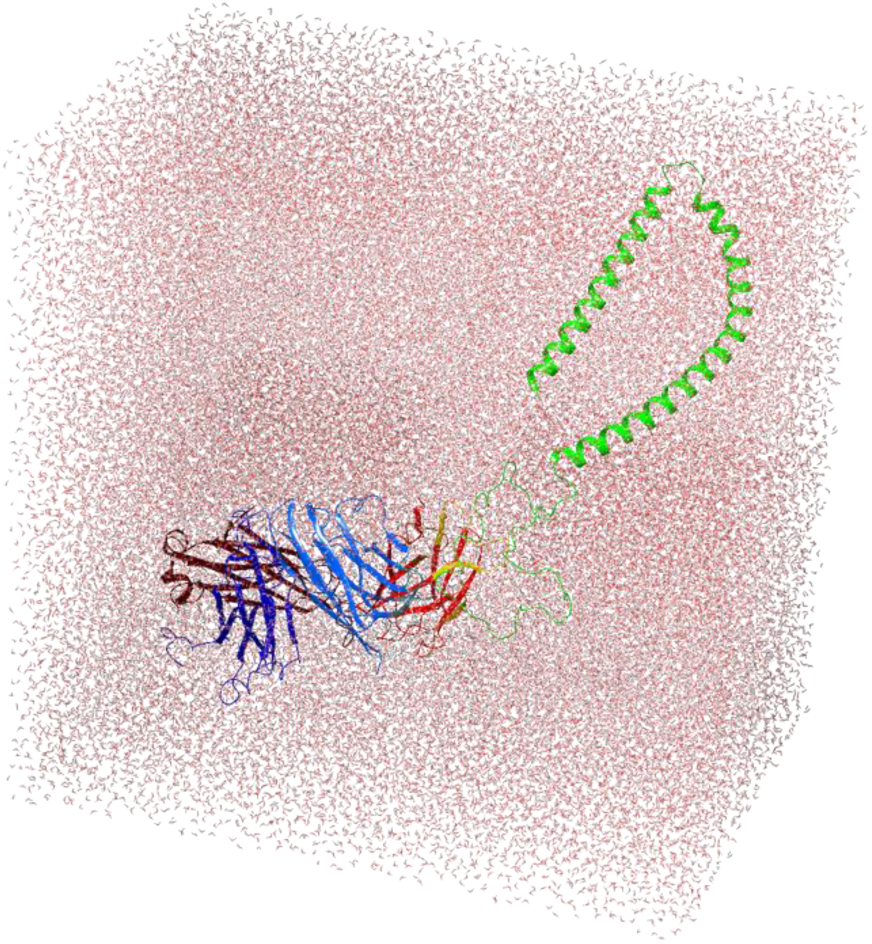
PRX002 and α-syn in a solvated and ionized system. The water and ion box extends 10 Å past the edge of the protein boundary defined by transferable intermolecular potential with the TIP3P water model.

### 2.3: Molecular Dynamics

All simulations were performed using VMD and NAMD (versions 2.14 and 3.0alpha). Simulations were run using A100 GPU on the Anvil high-performance computing cluster. Results from the ionization steps were minimized, heat prepared and brought to equilibrium before production runs were performed. Simulations were performed at 300 K, kept constant by Langevin dynamics, and pressure at 1.01325 Bar, kept constant by Langevin piston. After energy minimization was performed, the system was heated from 100 K to 300 K over 5 K intervals for 200 ps. The integration timestep of the simulation was 2 fs per step, with the position coordinates saved every 200 ps in a DCD file. Particle Mesh Ewald (PME) [46] were used for long-range electrostatic interactions, and a smooth cutoff was implemented for Van der Walls forces (10-12 Å).

### 2.4: MDAnalysis

Topology and coordinate files from the MD simulations were exported for descriptive data analysis of hydrogen bonds (h-bonds) formed during the simulation. MDAnalysis [47,48] was used to convert the coordinate files into NumPy [49] arrays and calculate the atoms on specific residues that were forming hydrogen bonds between the heavy and light chains of the PRX002 Fab and C terminus of α-syn (AA 100-140). Frames at which the h-bonds formed and the donor, hydrogen, and acceptor atom ID number. The number of h-bonds per timestep was calculated using the average bonds formed in each frame plotted using Matplotlib [50]. A reverse lookup function was utilized to list the residues forming hydrogen bonds, the chains where these residues were located, and the frequency of this unique bond’s appearance across all frames. PyMol was used to generate visualizations of the h-bonds formed between residues.

### 2.5: RMSD Analysis

To analyze the conformational changes of the V_H_ and V_L_ CDR regions, the root mean squared deviation (RMSD) was calculated. VMD’s RMSD analysis class was used to compute the difference between all frames of the simulation and the first frame. The PBC wrap function was used to ensure that the structure was centered in the simulation box before the RMSD calculation to prevent any periodic boundary condition-related errors.

### 2.6: Binding Free Energy Calculation

To measure the binding affinity of PRX002 with α-syn, gmx_MMPBSA was used for a generalized born and pairwise residue decomposition energy calculation [51]. Conversions of NAMD protein structure files and trajectories to GROMACS [52] format were performed using ParmEd [53] and MDTraj [54] respectively. The Generalized Born (GB) model was used to compute the predicted binding free energy by the following equation:

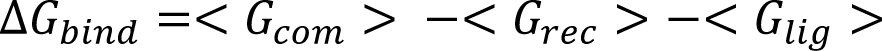

G_com_, G_rec_, and G_lig_ refer to the free energy of the complex, receptor, and ligand respectively. All 5000 frames of each 1 μs simulation were analyzed to determine their change in free energy.

### 2.7: Developability Methods

To determine the potential developability of PRX002, computational screening methods were used to determine in vitro viscosity, aggregation, and polyspecificity. DeepSCM was used to predict the spatial charge map (SCM) for screening potential viscosity issues. In addition, the Therapeutic Antibody Profiler (TAP) [55] was applied to screen other developability issues. These screening methods ensure the mAbs do not have elevated viscosity that would prevent subcutaneous administration and to see the similarity of the antibodies to clinical-stage biotherapeutics.

## 3. Results

### 3.1: Modelling of Antibody-Antigen Complex

Figure 2 shows the docked structures of the PRX002 Fab and α-syn. Docking simulations returned 10 best pose predictions, and the 3 outputs with the least distance between the CDR regions of the Fab with the C-terminus of α-syn were submitted for MD simulations. The three PRX002-α-syn complexes selected for MD simulations were 2KKW models 8, 14, and 27. (PD-8, PD-14, and PD-27). Chothia coloring schemes were utilized to highlight the specific CDR regions.

**Figure 2:**
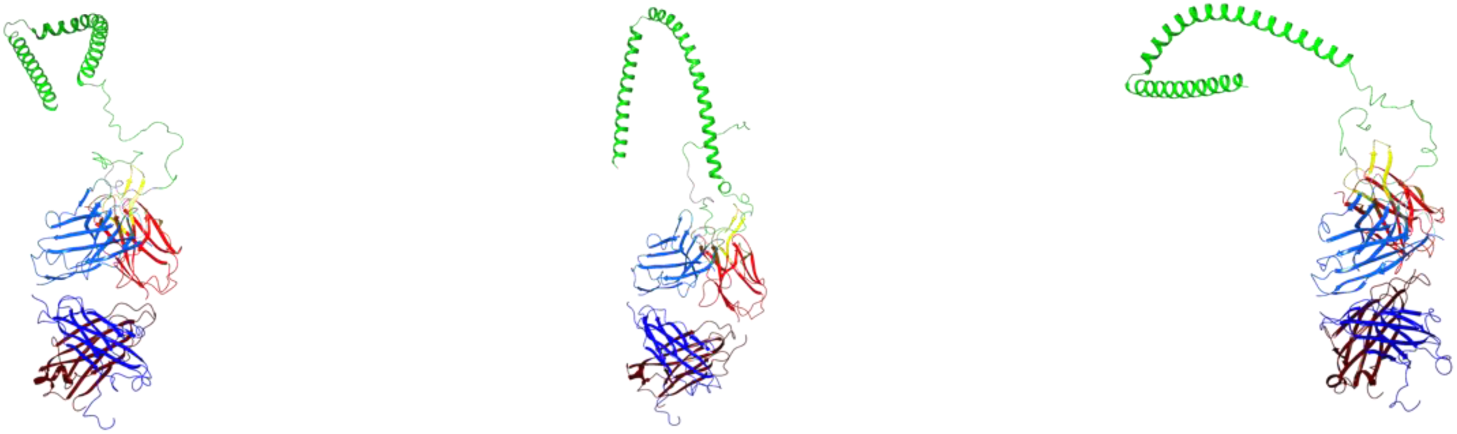
Complexes of PRX002 Fab region docked with α-syn. The heavy chain of PRX002 is highlighted in blue, the light chain of PRX002 is highlighted in red, and α-syn is highlighted in green. Protein-protein docking was performed by PIPER. The C-terminus (AA 118-126, highlighted in grey) of α-syn was specified in the protein-protein docking simulation as the attraction site. (From left to right: PD-8, PD-14, PD-27).

### 3.2: Hydrogen-Bond Analysis

Descriptive analysis of protein-protein h-bonds was performed on the outputs of the MD simulations to determine the location and frequency of intermolecular interactions over 1000 ns. Across the three separate simulations, an average of 6.368 h-bonds formed between the CDR regions of PRX002 and the C-terminus of α-syn. Tables 1-3 display the most prevalent bonds across the frames of the simulations. The percent occurrence of the h-bonds was calculated by taking the total amount of bonds formed and dividing it by the number of frames in the simulation. Some bonds, namely H:53-A:121 (Figure 3) in PD-8 and H:53-A:114 in PD-14 (Figure 4), formed more than one bond per frame, causing the percent occurrence to exceed 100%. For PD-27, the amount of hydrogen bonds was more evenly distributed in comparison to the previous two (Table 3), with the LCDR1 chain forming the most bonds (Figure 5). The second heavy chain CDR (HCDR2) region of PRX002 formed the most bonds with the C-terminus of α-syn.

**Table 1a-c:** A reverse lookup function was employed to take h-bond donor and acceptor id numbers to list the residues on each chain forming the most hydrogen bonds across the simulation. The top five unique residue pairs forming bonds were chosen. H = PRX002 Heavy Chain, L=PRX002 Light Chain, A=α-syn. Docked structures **(a)** PD8, **(b)** PD14, and **(c)** PD27.

System PD8
| H-Bond Donor | H-Bond Acceptor | Percent Occupancy |
| --- | --- | --- |
| Ser:H:53 | Asp:A:121 | 150.50% |
| Ser:H:52 | Asp:A:121 | 91.84% |
| Tyr:H:59 | Asp:A:121 | 90.30% |
| Tyr:L:38 | Glu:A:123 | 42.32% |
| Ser:L:99 | Tyr:A:125 | 28.94% |

System PD14
| H-Bond Donor | H-Bond Acceptor | Percent Occupancy |
| --- | --- | --- |
| Ser:H:53 | Glu:A:114 | 88.68% |
| Ser:H:52 | Glu:A:114 | 64.98% |
| Tyr:L:38 | Glu:A:111 | 58.76% |
| Tyr:H:59 | Glu:A:114 | 51.80% |
| Tyr:H:32 | Glu:A:110 | 50.56% |

System PD27
| H-Bond Donor | H-Bond Acceptor | Percent Occupancy |
| --- | --- | --- |
| Lys:L:36 | Asp:A:119 | 72.02% |
| Ser:L:66 | Glu:A:105 | 66.78% |
| Tyr:H:59 | Glu:A:126 | 58.96% |
| Tyr:L:38 | Glu:A:123 | 58.52% |
| Tyr:L:100 | Glu:A:126 | 55.48% |

**Table 2:** Therapeutic Antibody Profiler Results. Developability metrics provided from TAP analysis. CDR = Complementarity determining region, PSH = Patches of Surface Hydrophobicity, PPC = Patches of Positive Charge, PNC = Patches of Negative Charge, SFvCSP = Structural Fv Charge Symmetry Parameter

| Metric | Value |
| --- | --- |
| Total CDR Length | 49 |
| CDR Vicinity PSH | 147.4097 |
| CDR Vicinity PPC | 1.1106 |
| CDR Vicinity PNC | 0 |
| SFvCSP | -6 |

**Figures 3a-c:**
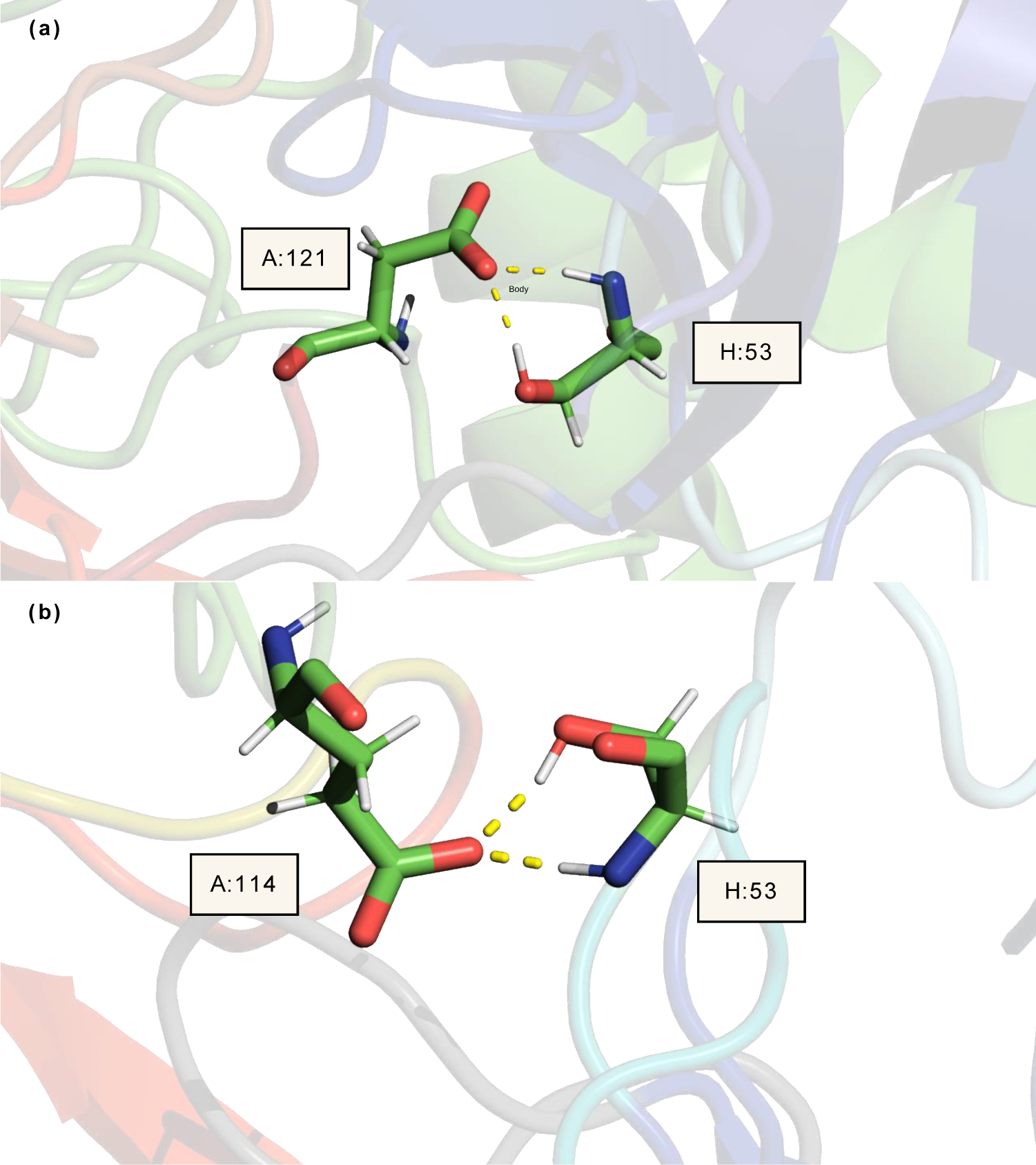
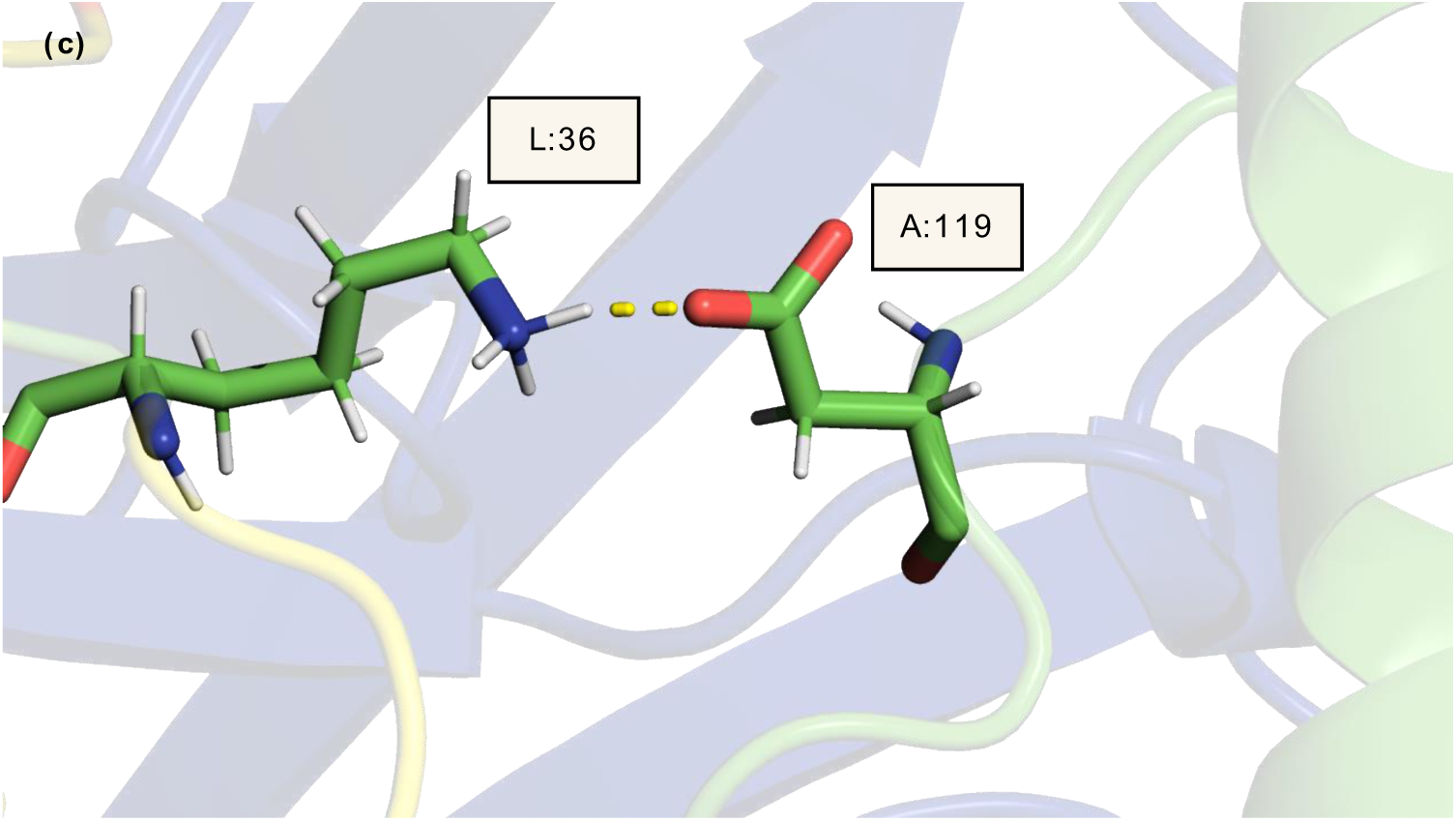
H-bonds formed between the variable region of PRX002 and α-syn. **(a)** Visualization of the h-bonds formed between serine on PRX002 HCDR2 region and aspartic acid on α-syn for system PD8. The NH1 and OH1 groups on serine are the donor atoms forming the bond with an OC group on aspartic acid. The most prevalent of these two interactions is Ser:OH1-Asp:OC. **(b)** Visualization of the h-bonds formed between serine on PRX002 HCDR2 region and glutamic acid on α-syn for system PD14. The NH1 and OH1 groups on serine are the donor atoms forming the bond with the OC group on glutamic acid. The most prevalent of these two interactions is Ser:OH1-Glu:OC. **(c)** Visualization of the h-bonds formed between lysine on PRX002 LCDR1 region and aspartic acid on α-syn for PD27. The NH3 group on lysine is the donor forming the bond with the OC group on aspartic acid. H = Heavy chain, L = Light chain, A = α-syn, Green = Carbon, Red = Oxygen, Blue = Nitrogen, White = Hydrogen

**Figures 4a-c:**
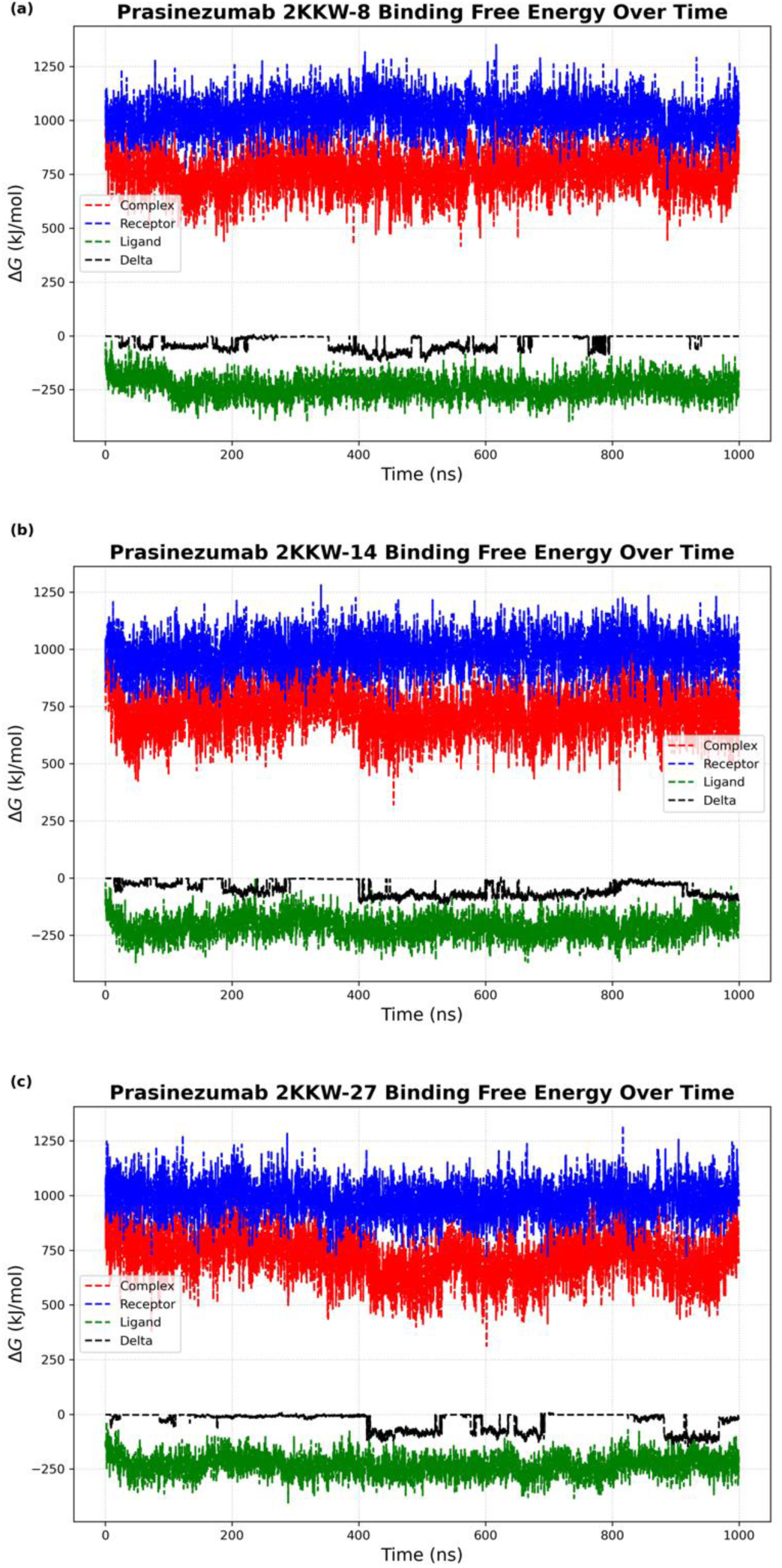
Analysis of predicted binding free energy estimates of PRX002 and a-syn simulations. All trajectories of systems **(a)** PD8, **(b)** PD14, and **(c)** PD27 were analyzed. Blue = Free energy of complex, Red = Free energy of receptor, Green = Free energy of ligand, Black = Total change in free energy of the system

**Figures 5a-c:**
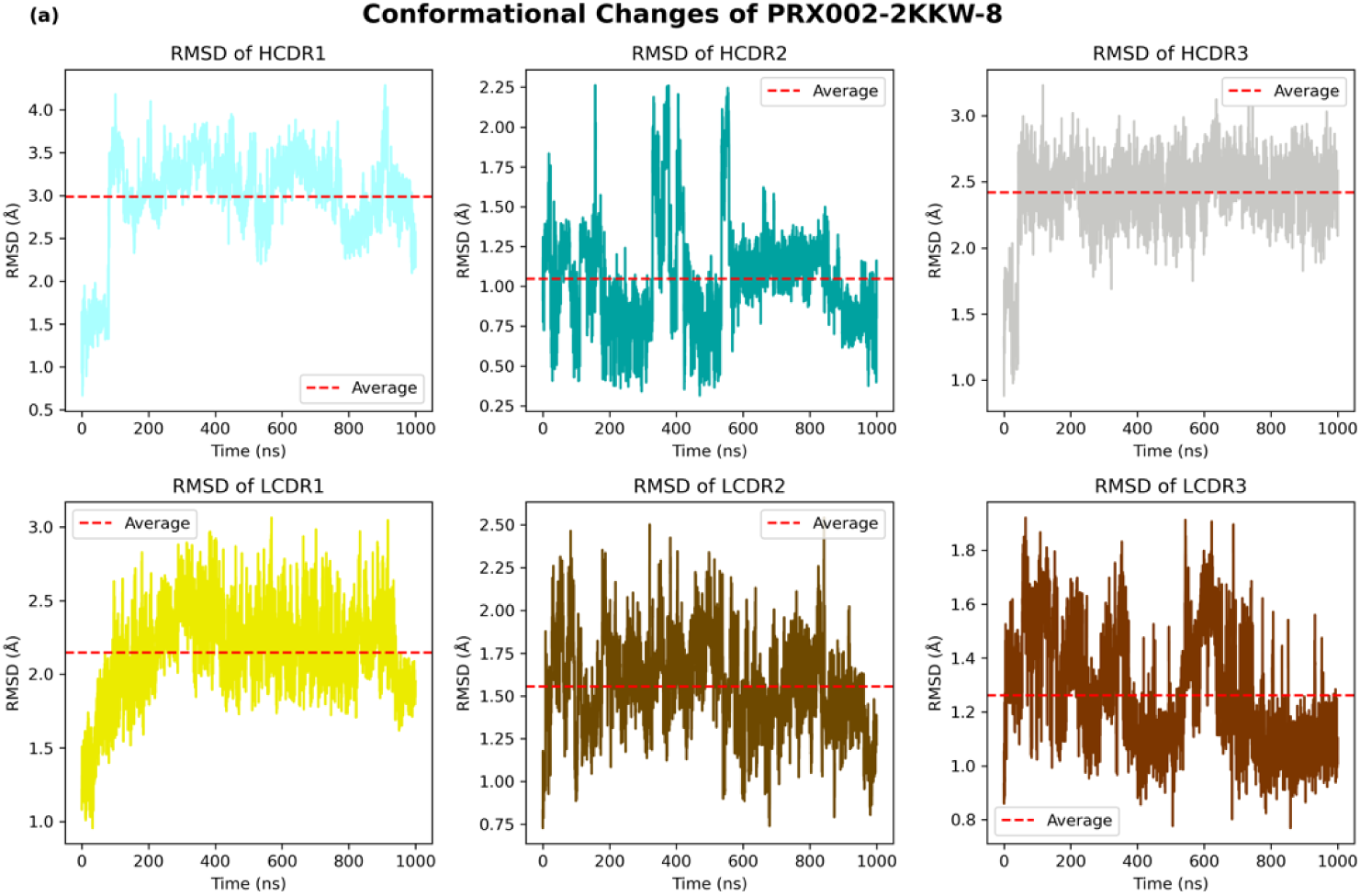
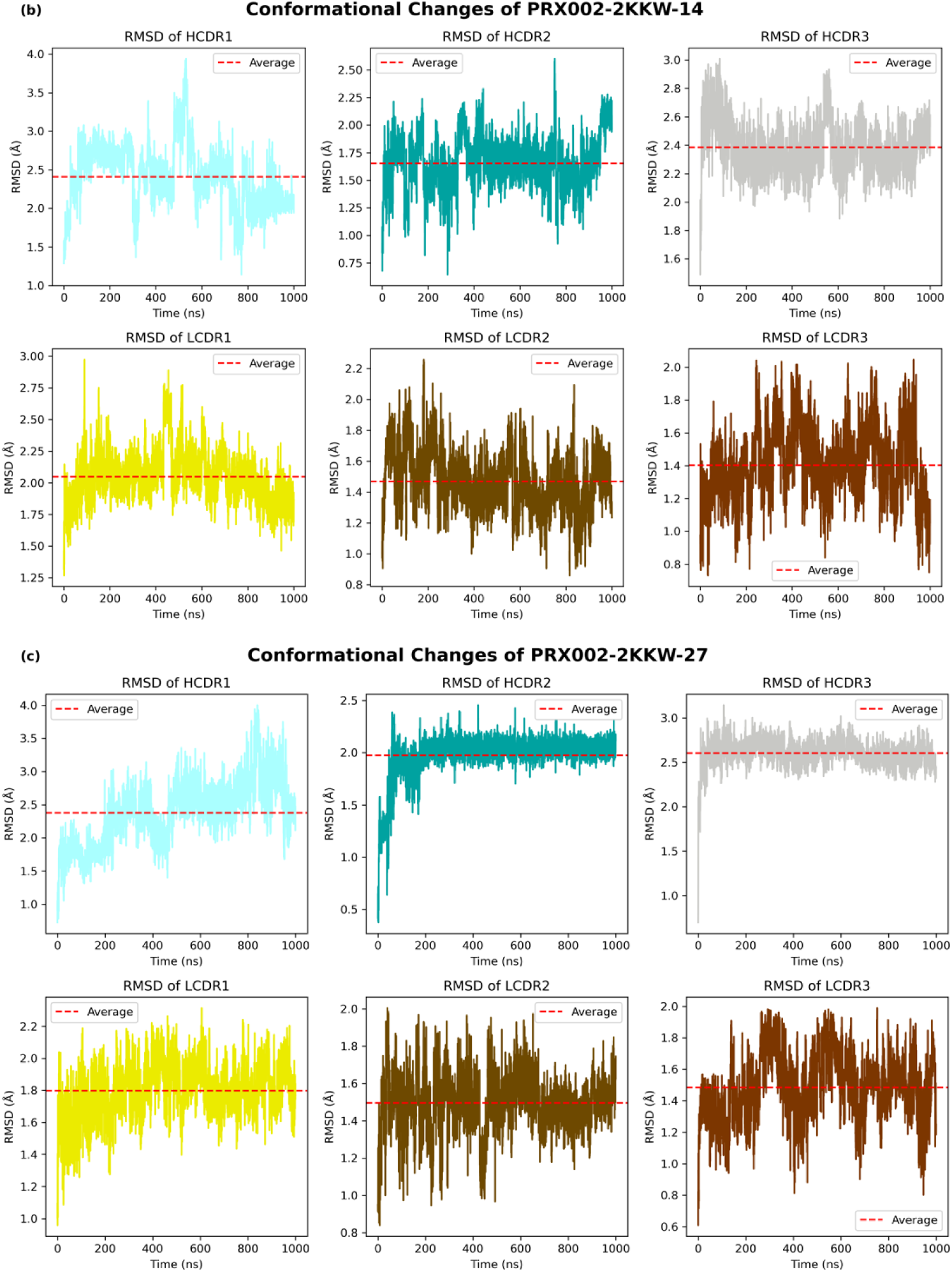
Conformational changes of CDR regions of PRX002 over the frames of simulations **(a)** PD-8, **(b)** PD14, and **(c)** PD27. The color of each subplot is analogous to the color of each CDR region defined by the Chothia numbering scheme.

### 3.3: Binding Affinity of Antibody-Antigen Complex

All 5000 frames of each of the 1 μs simulations were analyzed for free energy estimation by gmx_MMPBSA. The resultant average binding free energy for systems PD8, PD14, and PD27 was -28.3 -47.6 and -32.0 kJ/mol respectively. Change in binding free energy is shown in Figures 4a-c below.

### 3.4: Conformational Changes of CDR Regions During Binding

Figures 5a-c display the RMSD in angstroms (Å) of each CDR region of PRX002 over the frames of the simulation. For PD-8, the average RMSD for HCDR2 remains around 1.0 Å, with the lowest deviation compared to the rest of the CDR regions of the Fv region. The rest of the CDR regions do not exceed an average of 2.5 Å throughout simulation. In simulation PD-14, LCDR3 had the lowest average RMSD, at 1.4 Å, with LCDR2 following and HCDR2 after that, with RMSDs of 1.5 and 1.7 Å, respectively. The remainder of the values lie between 2.0-2.5 Å and relatively maintain consistent values throughout. PD-27 shows the highest RMSD values for the V_H_ CDR regions, with a 2.6 Å average for HCDR3, and LCDR2 having the lowest distribution at 1.5 Å.

### 3.5: Developability for Subcutaneous Administration

To test the developability of PRX002, DeepSCM was used to calculate the theoretical spatial charge map (SCM) score, with the mAb scoring 695.50, predicting low viscosity in high SC injection concentrations (the threshold value is 1000). TAP determined that the mAb was in range for all of the measured metrics except for SFvCSP, with it scoring -6.0, just on the boundary of the amber region, predicting that the antibody design of PRX002 is comparable to those of clinical-stage therapeutics.

## Discussion

Targeting aggregated α-syn has been a theorized route of treatment for years, but no medications have been developed to do so thus far. Prasinezumab is currently in Phase IIb and is the most promising candidate for a mAb targeting this pathologic protein. Understanding the mechanism by which the mAb binds to α-syn is vital for understanding its effectiveness and specific residues in the CDR regions that can be used for epitope mapping of future novel therapeutics. Our results show that the key intermolecular interaction of PRX002 with α-syn is Ser:H:53-Asp:A:118, evidence that the biotherapeutic does not separate from its target antigen and the stability of the CDR regions throughout the simulation. These findings indicate a plausible target in the C-terminus of α-syn for further epitope mapping and display the theoretical stability of PRX002 in human body conditions.

While Ser:H:53-Asp:A:118 is the most prevalent bond, evidence of hydrogen bonding between several residues on multiple CDR regions on both V_H_ and V_L_ chains is a sign of the strong intermolecular attraction driving the antibody-antigen recognition. The increased prevalence of binding residues on V_L_ CDR regions during PD-27 provides valuable insight into the behavior of both variable regions of PRX002 participating in binding. The estimated binding free energy calculated through the MM/PB(GB)SA calculation shows that PRX002 has a high predicted affinity for α-syn. Binding affinity is a fundamental metric in drug development and discovery, and showing a strong drive for antibody-antigen binding is vital for validating Prasinezumab as a potential therapeutic. Moreover, the procedures indicated in this paper provide a streamlined approach to assess the theoretical intermolecular attractions and binding affinity of a potential antibody therapeutic from sequence information alone, allowing for powerful insights early in the development process. Further experimental analysis to validate the intermolecular interactions between the mAb and α-syn would provide more evidence to support the ongoing clinical trial of Prasinezumab.

Several different tools were used for protein-protein docking of PRX002 and α-syn. ZDock [56], HDock [57], and PIPER were the rigid docking algorithm-based methods used to simulate the antibody-antigen interaction, with PIPER yielding the best results across all 34 variations of α-syn. PIPER being implemented in the BioLuminate platform also allows for easy preparation of proteins before simulation with other features, as well as assigning attraction and repulsion parameters for the docking algorithm. Generating structural fingerprints from the resultant protein-protein structure is also extremely valuable for feature generation, which can be implemented in machine learning algorithms.

The search to cure PD has spanned decades thus far and targeting α-syn has shown the greatest results both in the α-syn-pre-formed fibril model and human clinical trials. However, medications in the drug development pipeline have a high attrition rate, leading to millions of dollars put towards a medication that is either ineffective or undevelopable. Implementing computational tools such as MD for investigating the intermolecular forces, or lack thereof, will allow for quicker validation of a therapeutic agent or archival of an ineffectual candidate.

## Conclusion

Residues on the HCDR2 and LCDR1 of PRX002 form h-bonds with the C-terminus of α-syn throughout the MD simulations performed, and PRX002 has a strong affinity for α-syn. Our results show theoretically solid evidence of the binding capability of PRX002 in simulated human body conditions. PRX002 was also shown to not undergo drastic conformational changes after binding to α-syn, showing the strong lock-and-key recognition of this protein-protein complex. We propose that our method of calculating these intermolecular forces from the simulation will prove useful to future research of biopharmaceutical candidates for PD and beyond. In addition, we anticipate the usage of this methodology in early-stage development to determine the binding capabilities of biological drugs quickly.

## Data Availability

The code and scripts used in this study are freely available at https://github.com/malcolmharr/Mechanistic-study-for-Parkinsons-Disease.

## CRediT authorship contribution statement

**Malcolm Harrison:** Conceptualization, Methodology, Software, Data curation, Formal analysis, Visualization, Investigation, Writing – original draft.

**Pin-Kuang Lai:** Conceptualization, Methodology, Software, Supervision, Writing – review & editing.

## Acknowledgment

This work is supported by NSF REU/RET Site Grant #2050921. We acknowledge the computing resources provided by ACCESS (BIO220114 and CHM210013)

